# GhCPK28 negatively regulates the immune response by phosphorylating GhTIFY3b

**DOI:** 10.1101/2024.02.23.581816

**Authors:** Shengqi Gao, Wukui Shao, Jiawen Wu, Zhun Zhao, Wenran Hu, Panxia Shao, Jian Hua, Baohong Zou, Quansheng Huang

## Abstract

The soil-borne fungus *Verticillium dahliae* (*V. dahliae*) seriously inhibits cotton (*Gossypium hirsutum*) growth and productivity. The immune system of cotton against this pathogen is largely unknown. Here we investigated the involvement of Ca^2+^-dependent protein kinases (CPKs) in this immunity. One CPK coding gene *GhCPK28* had reduced expression after infection by *V. dahliae* in cotton. Knocking down of the *GhCPK28* by virus induced gene silencing led to enhanced resistance to *V. dahliae* which is accompanied by a higher ROS accumulation and systemic acquired response. GhCPK28 was found to phosphorylate a TIFY family protein GhTIFY3b and reduce its stability. Silencing *GhTIFY3b* increased the susceptibility of cotton to *V. dahliae*. Together, this study indicate that GhCPK28 is a negative regulator of defense against *V. dahliae* infection and the GhTIFY3b might be a target protein of its activity. It sheds light on the immune system against *V. dahliae* and provides candidate genes for improving resistance against Verticillium wilt in cotton.

## Introduction

Cotton (*Gossypium hirsutum*) is one of the world’s most important cash crops. However, the quality and yield of cotton is often seriously affected by Verticillium wilt caused by a soil-borne pathogen *Verticillium dahliae* (*V. dahliae*) (Atallah et al., 2011; Zhang et al., 2016). *V. dahliae* usually leads to vascular wilt through plant roots and forms a dormant structure of microsclerotia, which can survive for several years in soil without host and then infect subsequent crops (Shaban et al., 2018). Once plants are infected by *V. dahliae* it cannot been effectively control by fungicide. Although the selection and breeding of disease-resistant varieties are the fundamental measure for Verticillium wilt control, the key genes and regulatory mechanisms underlying cotton resistance to Verticillium wilt remain unclear (Song et al., 2020).

Plant immunity consists of two layers of defense responses including Pattern-triggered immunity (PTI) and effector-triggered immunity (ETI), each induced by a specific recognition of pathogen features by plant receptors (Jones and Dangl, 2006; Zhou and Zhang, 2020; Wang et al., 2022). PTI relies on the detection of microbe/pathogen-associated molecular patterns (MAMPs/PAMPs) outside the cell. MAMPs are usually defined as conserved features of microbes and damage-associated molecular patterns (DAMPs) are endogenous molecules released into the extracellular space due to damage caused by an invading pathogen (Bent and Mackey, 2007; Zipfel, 2014). Cell surface-localized pattern recognition receptors (PRRs) perceive conserved microbial MAMP or DAMP elicitors at the cell surface and transduce the signal inside the cell, triggering PTI (Dodds and Rathjen, 2010; Couto and Zipfel, 2016). PTI activates a few conserved downstream signaling pathways, including production of reactive oxygen species (ROS), activation of mitogen-activated protein kinases (MAPKs), and calcium influxes (Zipfel et al., 2006; Tian et al., 2019). Cytoplasmic Ca^2+^, as a second messenger, plays an important role in the infection of plant cells by pathogen (Gao et al., 2013). There are four major classes of Ca^2+^ sensors in plants such as calmodulins, calmodulin-like proteins, calcineurin B-like proteins, and Ca^2+^-dependent protein kinases (CPKs) (Bender and Snedden, 2013).

CPK can be activated by calcium signals during pathogen infection, inducing conformational change and activating kinase activity, leading to CPK itself phosphorylation and substrate phosphorylation (Cheng et al., 2002). For instance, the proteasome ATL31/6 targets AtCPK28 through degradation to ensure BIK1 homeostasis and dynamically regulate immune signaling (Liu et al., 2021). Silencing *NaCPK4* and *NaCPK5* genes, homologs of *AtCPK28, attenuate* led to accumulation of many defensive metabolites are accumulated in tobacco (Yang et al., 2012). Loss of function of *OsCPK4* and *OsCPK18* homologous to *AtCPK28* in rice shows enhanced immune signal transduction and resistance to pathogens infection (Monaghan, 2018). Additionally, rice *OsCPK9* plays an important role in the signal transduction of rice blast response (Asano et al., 2005). *TaCPK2*, a calcium-dependent protein kinase gene that is required for wheat powdery mildew resistance enhances bacterial blight resistance in transgenic rice (Geng et al., 2013). *TaCPK7* positively regulates the wheat resistance response to *Rhizoctonia cerealis* infection through the modulation of the expression of defence-associated gene (Wei et al., 2016). *CaCPK15* indirectly activates downstream *CaWRKY40* which in turn potentiates and increases the susceptibility of pepper to *Ralstonia solanacearum* inoculation (Shen et al., 2016). These reports demonstrate that *CPK* protein participates in plant defense response by regulating reactive oxygen species accumulation and defense-related phytohormone biosynthesis.

Hormone-mediated signaling is one of the important defense mechanisms of cotton against Verticillium wilt. Among them, jasmonate (JA), salicylic acid (SA), and ethylene (ET) play important roles in the stress signaling pathway of plants infected with pathogens (Sun et al., 2014; Zhang et al., 2017). After plant infection with pathogens, the immunity induced by PTI and ETI effects usually in a short period of time regulated JA synthesis and increased JA content (Fradin et al., 2011). JA synthesis-related genes, such as JAZ (Jasmonic acid ZIM domain protein) and AOS (allene oxide synthase), dynamically regulate JA synthesis after pathogen infection. JA plays an important role in enhancing the resistance of *Arabidopsis* to *V. dahliae* and other fungi (Thatcher et al., 2016). The TIFY gene family is plant specific and encodes putative transcription factors (TFs) characterized by a highly conserved motif (TIF[F/Y]XG) positioned within an approximately 28-amino-acid TIFY domain (Vanholme et al., 2007; Bai et al., 2011). Based on phylogenetic and structural analyses, genes of this family can be divided into four groups, viz. ZML(ZIM-like), TIFY, PPD(PEAPOD), and JASMONATE ZIM (ZINC-FINGER EXPRESSED IN INFLORESCENCE MERISTEM)-domain (JAZ) proteins. TIFY family members, particularly JAZ subfamily proteins, play roles in biological processes such as stress and hormone responses. JAZ subfamily members are involved in the jasmonic acid (JA) pathway (Yan et al., 2007). *AtTIFY10b*/*JAZ1* is an important player in JA signal transduction. To stop their repression of JA-response genes, *JAZ1* is degraded in the presence of JA through the SCF^COI1^-dependent 26S proteasome pathway (Thines et al., 2007). JAZ proteins in the absence of JA interact with and repress the basic helix-loop-helix (bHLH) TFs MYC2 and MYC3. These TFs bind directly to DNA sequences and regulate the expression of downstream JA-responsive genes (Chini et al., 2007). *JAZ* genes also show differential induction in response to JA treatment, herbivory, wounding, *Pseudomonas syringae* infection, and environmental stresses such as drought, low temperature and salinity (Chung et al., 2008; Ye et al., 2009; Demianski et al., 2011)

There are six copies (Gh_A02G1635, Gh_D03G0087, Gh_A10G0886, Gh_D10G0863, Gh_D11G1774 and Gh_A11G1615) of *GhCPK28* in the cotton genome. Previous studies have shown that these *GhCPK28* have different expression patterns under *V. dahliae* stress. Gh_D10G0863 and Gh_A10G0886 were down-regulated while Gh_D11G1774 and Gh_A11G1615 were up-regulated. Gh_D03G0087 had no significant change. The knockdown of *GhCPK28* (Gh_A11G1615) expression led to the increase of reactive oxygen species in cotton plants, a series of defense responses were enhanced, and the sensitivity of cotton to *V. dahliae* was decreased (Wu et al., 2021). We hypothesized that *GhCPK28* homologues may have a conserved function in negatively regulate immunity in various plant species. In this study, we identified a *CPK28* gene from upland cotton, *GhCPK28* (Gh_A10G0886), that is involved in regulating host defense against *V. dahliae* infection. Silencing *GhCPK28* expression enhanced the disease resistance of cotton plants against *V. dahliae* infection. Further investigation revealed that GhCPK28 interacts with and phosphorylates a TIFY family protein GhTIFY3b. Our results indicate that GhCPK28 is a negative regulator of defense against *V. dahliae* infection and the GhTIFY3b might be a target protein of its activity. It sheds light on the immune system against *V. dahliae* and provides candidate genes for improving resistance against Verticillium wilt in cotton.

## Results

### Expression profile of *GhCPK28* gene

Due to *CPK* is important in plant immunity, we analyzed the expression of all putative *CPK* genes in cotton with derived from cotton cv TM-1 roots infected by the *V. dahliae* isolate V991. Among these *CPK* genes, we found that a sequence (Gh_A10G0886) was down-regulated in the cotton root infected by *V. dahliae* strain V991 (Fig. 1A). RT-qPCR analysis confirmed that the *GhCPK28* was induced at 3 hpi in roots treated with H_2_O_2_ and JA, while there was no significant change in SA treatment compared with control (Fig. 1B) and was expressed in several organs (Fig. 1C). This finding indicated that the *GhCPK28* may be involved in responses to *V. dahliae* infection, ROS and JA.

**Figure 1.**
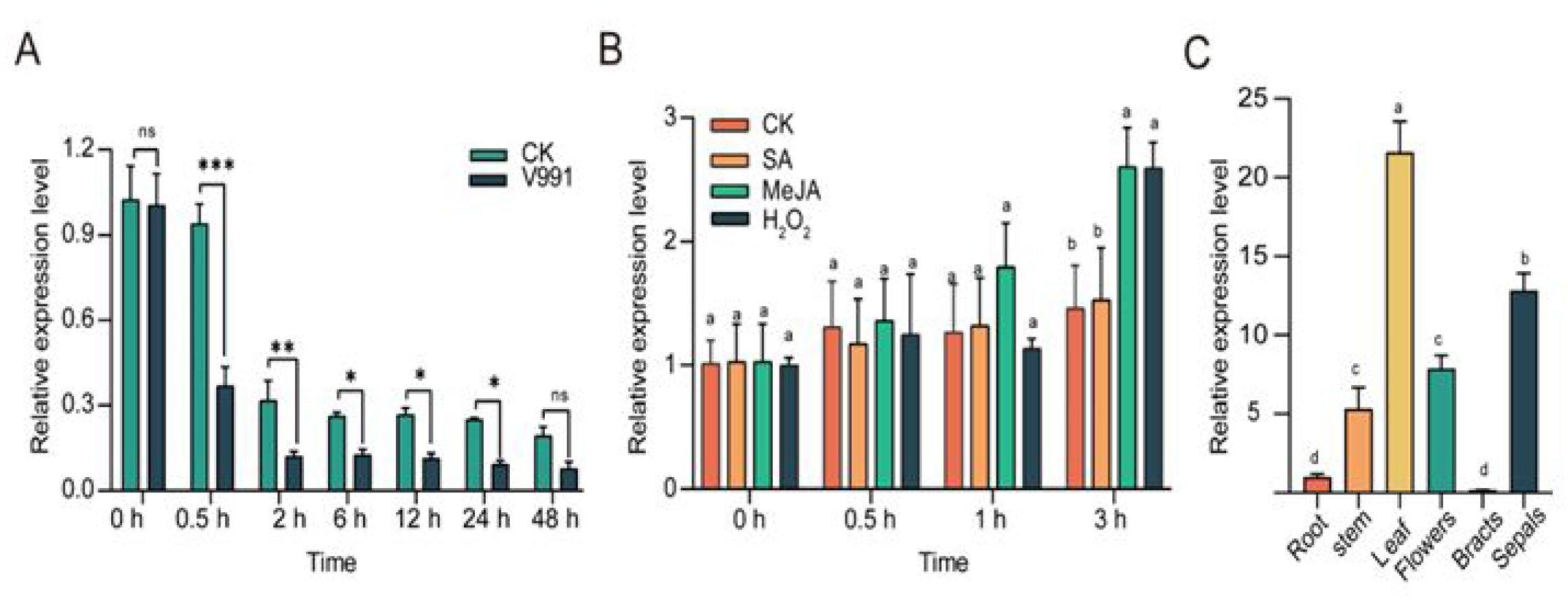
Expression pattern of *GhCPK28*. A, qRT-PCR time-course analysis of *GhCPK28* expression in response to *V. dahliae* infection. Error bars represent ± SD (n=3). The standard deviations were calculated from the results of three independent experiments (* P< 0.05, ** P< 0.01, t-test). B, qRT-PCR time-course analysis of *GhCPK28* expression in plants treated with JA, SA and H_2_O_2_. C, qRT-PCR analysis of *GhCPK28* expression in roots, stems, leaf, flowers, bracts and sepals of TM-1. h means hours post inoculation. *GhUBQ7* was used as the control. The values are means±sd, n=3. Different letters indicate significant differences at P< 0.05(ANOVA’s multiple comparison test).

The full-length cDNA of the sequence is composed of 1635 nucleotides encoding 544 amino acids, including N-terminal myristoylation sequence and palmitoylation site. The protein contains four characteristic domains: the N-terminal variable region, the Ser/Thr kinase catalytic domain, the autoregulatory/autoinhibitory domain and four conserved EF hands for Ca^2+^ binding (Supplemental Fig. S1A). Phylogenetic analysis of the protein sequence showed that the closest homologous gene to the protein was *AtCPK28* (*Arabidopsis thaliana*), *SlCPK28* (*Solanum lycopersicum*), *GrCPK28* (*Gossypium raimondii*), *ZmCPK28* (*Zea may*), and *OsCPK28* (*Oryza sativa*) (Supplemental Fig.S1B). Therefore, the gene was designated *GhCPK28*.

### Silencing *GhCPK28* leads to enhanced resistance to *V. dahliae*

To elucidate the function of *GhCPK28* in cotton resistance to Verticillium wilt, virus induced gene silencing (VIGS) technique was used to silence *GhCPK28* gene. Plants with the control gene *Cloroplastos alterados 1* (*CLA1*) silenced by *TRV:GhCLA1* showed bleaching phenotypes in new true leaves, showing that the VIGS experiments were effective (Fig. 2A). RT-qPCR analysis showed that the expression of *GhCPK28* was significantly decreased in *TRV:GhCPK28* silenced cotton plants (Fig.2B). The *TRV:GhCPK28* and *TRV:00* plants were inoculated with *V. dahliae* V991. Control plants exhibited more severe disease symptoms than those of *GhCPK28* silenced plants at 16 days (Fig. 2C). The rate of diseased plants and disease index of *TRV:GhCPK28* silent plants were both significantly lower than that of control *TRV:00* plants (Fig. 2D, E). Meanwhile, the stem segments of cotton plants silenced by *TRV:GhCPK28* had lower reproductive number of pathogens in stem segments compared to the control plants (Fig. 2F). The change of vascular bundle brown in control plants was more obvious than that in *GhCPK28* silent plants (Fig. 2G). Less fungal DNA was also observed in *GhCPK28*-silenced plants by fungal DNA abundance detection (Fig. 2H). These results suggest that *GhCPK28*-silenced cotton plants increases the resistance to Verticillium wilt, and *GhCPK28* is a negative regulator of Verticillium wilt in cotton.

**Figure 2.**
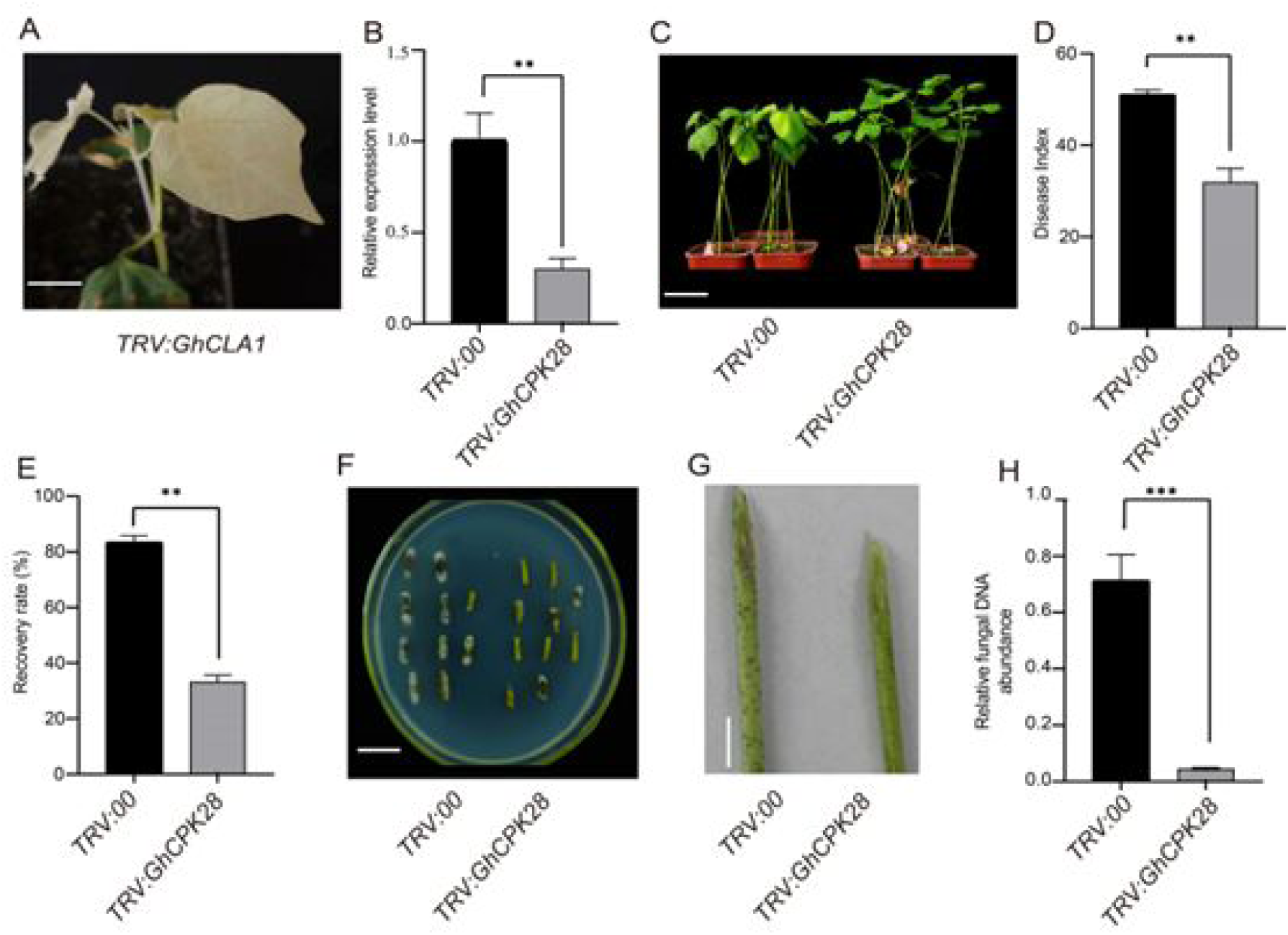
Silencing *GhCPK28* leads to enhance *V. dahliae* resistance. A, Photobleaching phenotype of cotton seedlings was observed after 16 days infiltrated with *TRV:GhCLA1*. B, qRT-PCR analysis of *GhCPK28* expression after inoculated with *TRV:GhCPK28*. C, Disease symptoms of control (*TRV:00*) and *GhCPK28*-silenced (*TRV:GhCPK28*) plants after inoculated with *V. dahliae*. D, Disease index of *TRV:00* and *TRV:GhCPK28* plants. E, Rate of diseased on *TRV:00* and *TRV:GhCPK28* cotton plants. F, Cotton stem segments 20 d after *V. dahliae* infection were plated on potato dextrose agar medium. Photos were taken 6 d after culturing. G, Comparison of darkened vascular tissues at 20 d after *V. dahliae* infection between *TRV:00* and *TRV:GhCPK28*. H, Fungal DNA relative amount detection. The total DNA of cotton stems 20 d after *V. dahliae* infection was extracted and used as a template for the detection of fungal biomass by RT-qPCR. *GhUBQ7* was used as control. Error bars represent ±SD (n=3). The standard deviations were calculated from the results of three independent experiments (* P< 0.05, ** P< 0.01, t-test).

### Downregulation of *GhCPK28* leads to increased ROS accumulation and defense response genes

Given that *GhCPK28* functions as a negative regulator against *V. dahliae* infection and could be induced by H_2_O_2_ stress, we speculate that the increased resistance to *V. dahliae* infection in *GhCPK28*-silenced plants may lead to the changes of the redox status. To determine this possibility, the production of total ROS was examined in the leaves of *GhCPK28*-silenced and control plants after *V. dahliae* infection. *GhCPK28*-silenced leaves were observed to accumulate more ROS than the control with DAB (3,3’-diaminobenzidine tetrahydrochloride) staining (Fig. 3A). Trypan blue staining revealed that there was less cell death in GhCPK28-silenced leaves than the control plants (Fig. 3B). This finding indicated that silencing *GhCPK28* inhibited cell death but increased ROS and confirmed that *GhCPK28* played a role in ROS burst that occurred challenging *V. dahliae* attack, thereby inducing less cell death.

**Figure 3.**
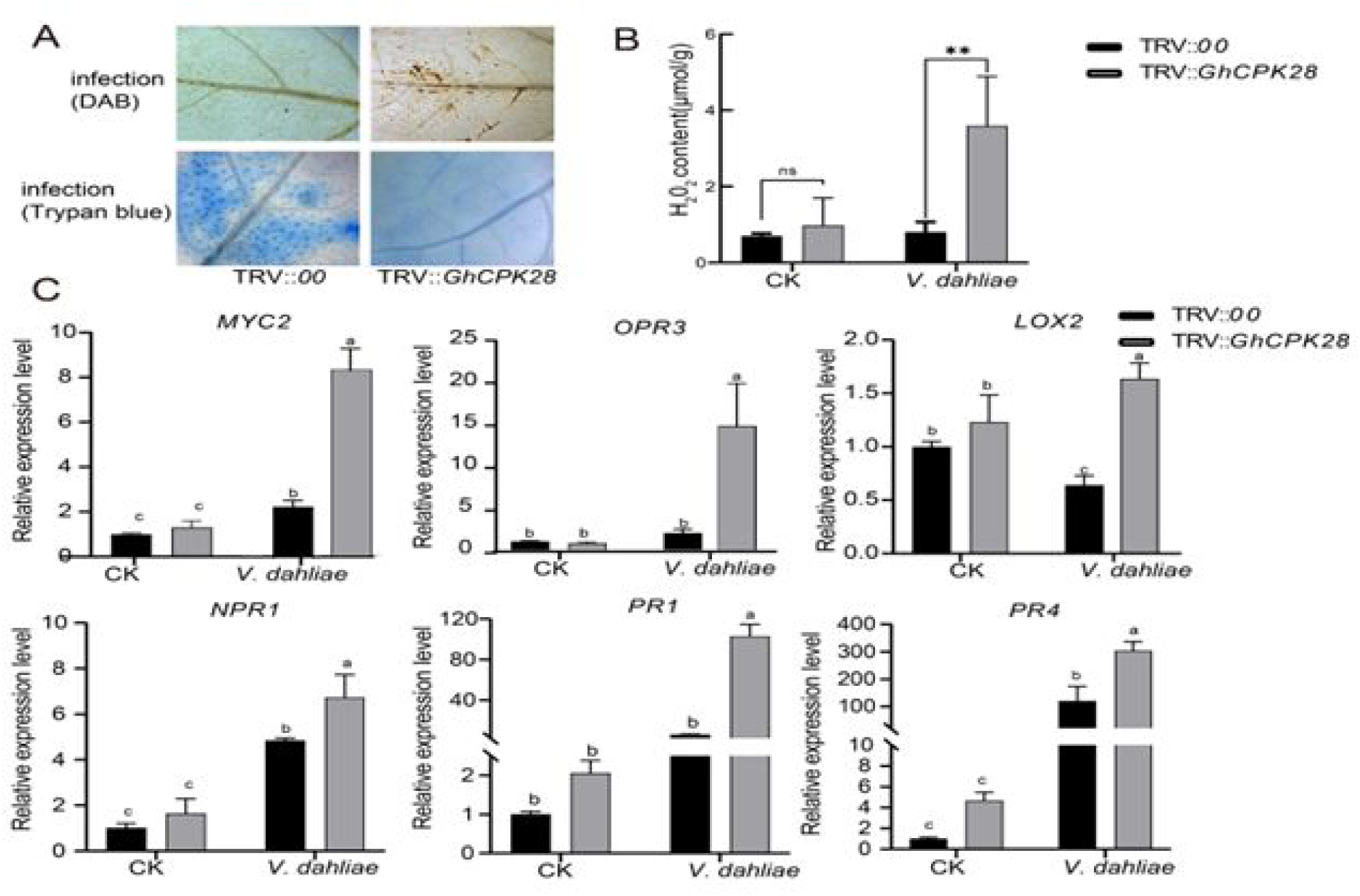
Silencing of *GhCPK28* results in ROS elimination and defense response to *V. dahliae.* A, Observation of ROS by DAB-peroxidase staining and cell death by Trypan blue staining 24 h after inoculation with *V. dahliae* in *TRV:00* and *TRV:GhCPK28* plants. B, H_2_O_2_ content in roots of *TRV:00* and *TRV:GhCPK28* plants 1 h after inoculation with *V. dahliae.* C, qRT-PCR analysis of pathogenesis and JA synthesis-related genes in *TRV:00* and *TRV:GhCPK28* cotton plants. Error bars represent SD of three technical replicates of one biological experiment. Different letters indicate significant differences at P< 0.05(ANOVA’s multiple comparison test).

Activation of plant defense responses during *V. dahliae* infection is accompanied by increase of jasmonate (JA) and the up-regulation of pathogenesis-related (PR) genes. To further characterize the disease resistance phenotype, the expression of PR genes such as *GhNPR1*(*Nonexpresser of Pathogenesis Related 1*), *GhPR1* (*Pathogenesis Related 1*) and *GhPR4* (*Pathogenesis Related 4*) and JA synthesis related gene *GhMYC2* (Myelocyto-matosis), *GhOPR3* (12-oxophytodienoate reductase) and *GhLOX2* (Lipoxygenases) was analyzed in *TRV:GhCPK28* and *TRV:00* plants after *V. dahliae* infection. As expected, *V. dahliae* challenge strongly induced the expression of the above genes in *TRV:00* plants. In *TRV:GhCPK28* plants, *GhPR1, GhNPR1* and *GhPR4* were induced with higher levels than in the wild-type plants. While the induced expression of *GhMYC2*, *GhOPR3* and *GhLOX2* were significantly increased in *TRV:GhCPK28* plants after *V. dahliae* (Fig. 3C). This result implied that the *GhCPK28* might regulate cotton immunity through modifying JA defense signaling pathways.

### GhCPK28 interacts with GhTIFY3b

To elucidate the regulatory function of *GhCPK28* in disease resistance in cotton, full-length *GhCPK28* was used to identify potential interacting proteins using a yeast two-hybrid (Y2H) library of cotton. Forty-nine candidates GhCPK28-interacting proteins were found in the Y2H screening (Supplemental Table S2). We further tested direct interaction between GhCPK28 and its candidate interacting proteins with a focus on GhTIFY3b (Gh_D05G039400.1). GhTIFY3b showed 50% identity compared with AtTIFY3b amino acid sequence (Supplemental Fig. S2). To confirm this interaction, the full-length *GhCPK28* and *GhTIFY3b* were constructed and transformed into the expressed yeast cells. It was found that GhCPK28 could interact with GhTIFY3b in the yeast two-hybrid assay (Fig. 4A). We further tested the interaction between GhCPK28 and GhTIFY3b by in vitro pull-down assays. GhCPK28 fused to His fusion recombinant protein (His) and GhTIFY3b fused to glutathione S-transferase (GST) were expressed in *E. coli*. The recombinant GhTIFY3b protein was able to pull down the GhCPK28 protein, but not His alone. Additionally, GST did not pull down the GhCPK28-His protein (Fig. 4B). These data indicate that GhCPK28 and GhTIFY3b proteins interact physically in vitro. The physical interactions between GhCPK28 and GhTIFY3b were further supported by assays of BiFC and co-immunoprecipitation (Co-IP). In BiFC, GhCPK28 protein and GhTIFY3b were respectively fused to the C-terminal half and N-terminal half of the YFP protein. Co-expression of GhCPK28-nYFP with GhTIFY3b-cYFP, but not with -cYFP, generated positive signals when co-expressed in *Nicotiana benthamiana* (Fig. 4C). In contrast, co-expression of nYFP with GhTIFY3b-cYFP did not yield positive signals. For Co-IP, flag tagged GhCPK28 was co-expressed with MYC tagged GhTIFY3b in *N. benthamiana*. GhTIFY3b was detected in fractions IPed by anti-Myc antibody when co-expressed with GhCPK28-flag but not with Myc tag alone (Fig. 4D). These results indicate that GhCPK28 can have direct physical interactions with GhTIFY3b.

**Figure 4.**
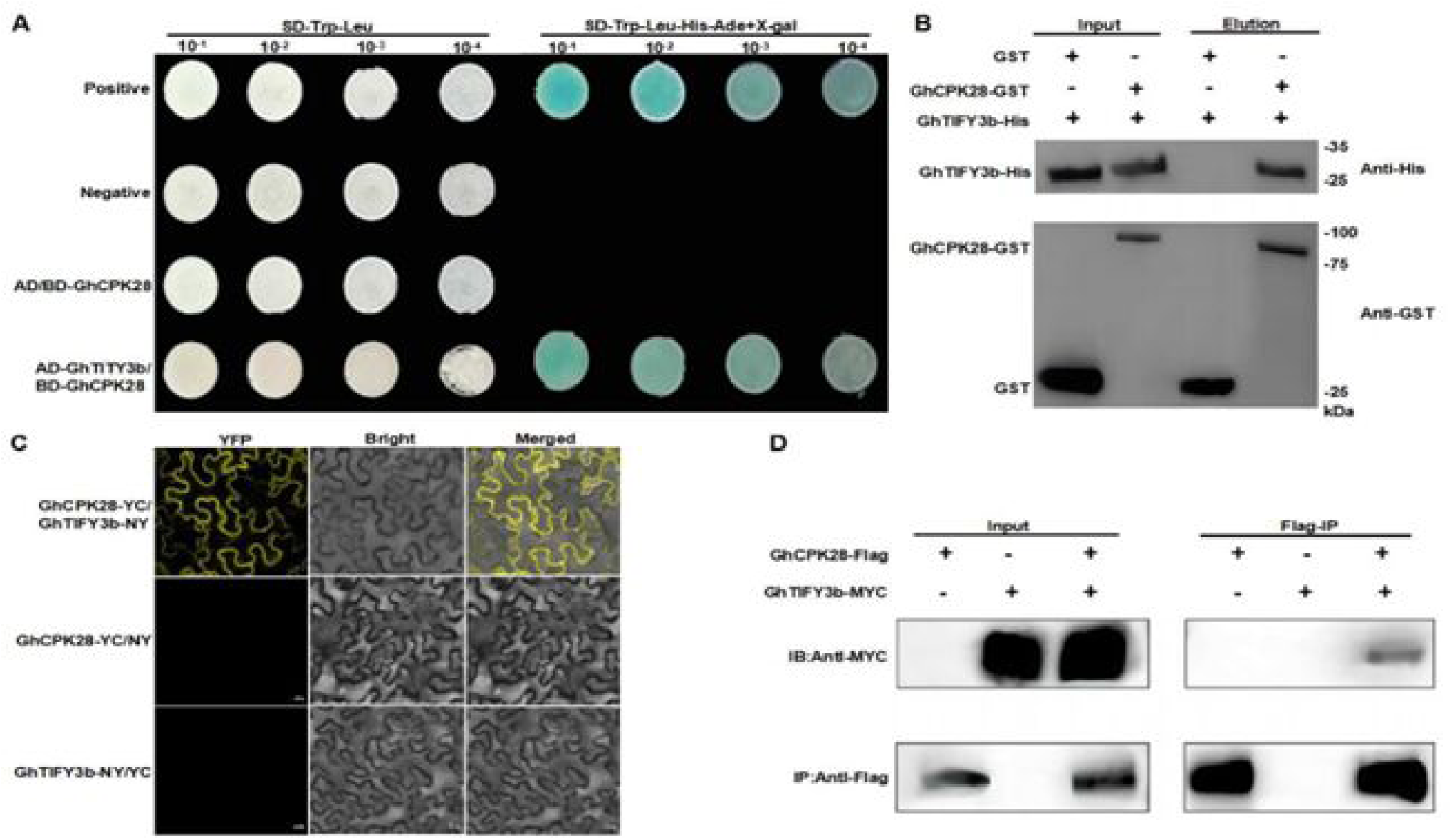
GhCPK28 interacts with GhTIFY3b in vitro and in vivo. A, Y2H assays showed the interaction between GhCPK28 and GhTIFY3b. The transformed yeast cells were grown on a synthetic dextrose (SD) media. Blue colonies on SD-Trp-Leu-His-Ade (containing X-α-Gal) exhibit positive interactions. B, GST pull-down assay showed direct interactions between GST-GhCPK28 and His-GhTIFY3 fusion proteins. His-GhTIFY3b protein was incubated with immobilized GST or GST-GhCPK28 protein, and the immunoprecipitation portion was detected with anti-His antibody or anti-GST antibody. C, BiFC assay showed that GhCPK28-cYFP interacted with GhTIFY3b-nYFP to form functional YFP on the plasma membrane. The interaction of GhCPK28-cYFP with GhTIFY3b-nYFP, GhCPK28-cYFP with nYFP, GhTIFY3b-nYFP with cYFP were used to detect BiFC. Merged = merging of YFP autofluorescence. D, Co-IP assay showed that GhCPK28 interacted directly with GhTIFY3b in vivo. The clones carrying GhCPK28-flag and GhTIFY3b-MYC were coinfiltrated into *N. benthamiana* leaves by *Agrobacterium tumefaciens*. Total protein was immunoprecipitated with anti-flag and anti-Myc antibodies, respectively. All the experiments were repeated at least three times with similar results. Bars = 20 μm.

Additionally, we constructed GhCPK28-GFP and GhTIFY3b-GFP fusion expression vectors and expressed them transiently in *N. benthamiana* leaves, respectively. GhCPK28-GFP was detected mainly in cell membrane, and GhTIFY3b-GFP was observed only in nucleus (Supplemental Fig. S3).

### Silencing of *GhTIFY3b* in cotton increases susceptibility to *V. dahliae*

Since GhCPK28 can interact with GhTIFY3b, we tested whether GhTIFY3b regulated cotton resistance to *V. dahliae*. *GhTIFY3b* expression was induced by *V. dahliae* (Supplemental Fig. S4A) but down regulated by SA, H_2_O_2_ and JA treatment (Supplemental Fig. S4B). The *G. hirsutum* genome contains six homologous *TIFY3b* genes: Gh_A05G029600, Gh_D05G039400, Gh_A07G018400, Gh_D07G019800, Gh_A09G200600 and Gh_D09G193800. Like other TIFY3b proteins, GhTIFY3b contains a highly conserved motif (TIF[F/Y]XG) (Supplemental Fig. S5).

We used VIGS to study the effect *GhTIFY3b* (Gh_D05G039400) on resistance of cotton resistance to *V. dahliae*. Plants inoculated with *TRV:GhCLA1* served as positive control for VIGS efficiency (Fig. 5A). *GhTIFY3b* expression level was significantly reduced in *TRV:GhTIFY3b*, but not in *TRV:00* control plants (Fig. 5B). After *V. dahliae* infection, the disease symptoms such as leaf wilting and yellowing of *TRV:GhTIFY3b* plants were more serious than those of *TRV:00* plants (Fig. 5C), resulting in significantly higher disease index (Fig. 5D). The fungal recovery test using stems of inoculated cotton plants further demonstrated a higher level of *V. dahliae* colonization in *TRV:GhTIFY3b* than in *TRV:00* plants (Fig. 5E) with significant more browning of vascular tissue (Fig. 5F). In addition, ROS levels in *TRV:00* leaves were higher than *TRV:GhTIFY3b* leaves 24 hours after inoculation with *V. dahliae*, and *TRV:GhTIFY3b* leaves had more cells death in leaves than *TRV:00* leaves (Supplemental Fig. S6A). Down-regulation of *GhTIFY3b* appears to have a negative effect on NO accumulation in *V. dahliae* infected plants (Supplemental Fig.S6B). These results clearly showed that knockdown of *GhTIFY3b* weakened the resistance of cotton to *V. dahliae*.

**Figure 5.**
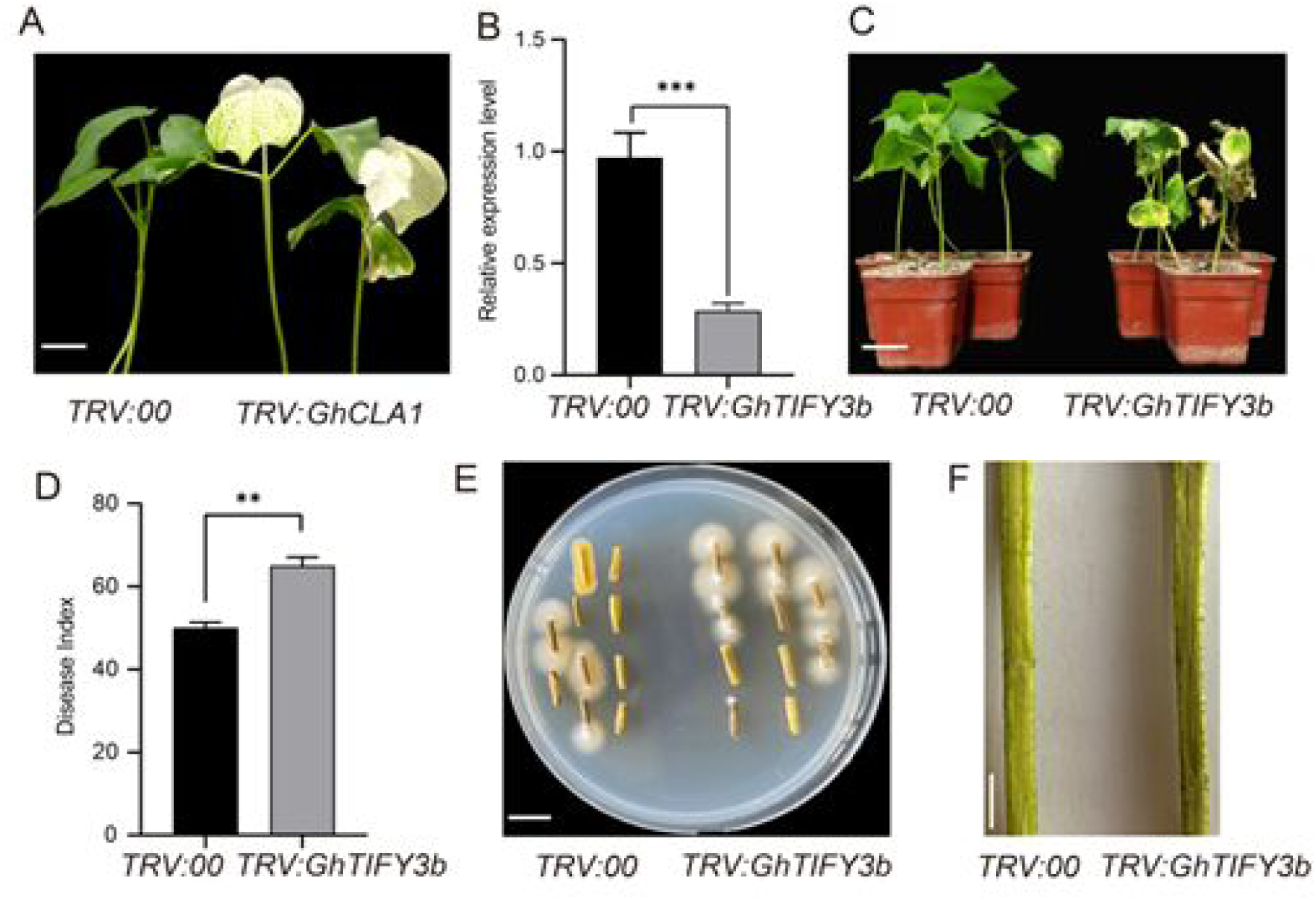
Silencing *GhTIFY3b* leads to increase *V. dahliae* susceptibility. A, Photobleaching phenotype of *TRV:GhCLA1* plants at 16 days. B, *GhTIFY3b* expression level by qRT-PCR analysis in plants of *TRV:GhTIFY3b*. C, Disease symptom after inoculation with *V. dahliae* in control (*TRV:00*) and *GhTIFY3b*-silenced (*TRV:GhTIFY3b*) plants. D, Disease index of *TRV:00* and *TRV:GhTIFY3b* plants. E, Cotton stem segments 20 d after infection with *V. dahliae* were plated on PDA medium. Photos were taken 6 d after culturing. F, Comparison of darkened vascular tissues at 20 d after infection with *V. dahliae* between *TRV:00* and *TRV:GhCPK28*. Error bars represent ±SD (n=3). The standard deviations were calculated from the results of three independent experiments (* P< 0.05, ** P< 0.01, t-test). Bars =2 cm.

### GhCPK28 direct phosphorylates GhTIFY3b

In order to determine whether GhCPK28 phosphorylates GhTIFY3b, we conducted in vitro phosphorylation analysis of GST-GhCPK28 and His-GhTIFY3b, and radio-autograph showed that GhCPK28 could phosphorylate GhTIFY3b. Moreover, the phosphorylation degree enhanced with the increase of the concentration of the two proteins (Fig. 6A). Subsequently, liquid chromatography-tandem mass spectrometry was applied to identify the probable GhCPK28 phosphorylation sites in GhTIFY3b. The Thr-110 was identified as one of the most abundant phosphorylation sites in GhTIFY3b after incubating with GhCPK28 (Supplemental Fig. S7). To further verify the phosphorylation site of GhTIFY3b, the Thr-110 site of GhTIFY3b was mutated to Ala (GhTIFY3b-mA) or Asp (GhTIFY3b-mD). His-tagged phosphodeficient GhTIFY3b-mA, phosphomimic GhTIFY3b-mD and GhCPK28-GST recombinant proteins were generated and respectively subjected to GST Pull-Down experiment. His-GhTIFY3b-mA and His-GhTIFY3b-mD had the same physical interactions with GhCPK28 as the wild-type GhTIFY3n (Supplemental Fig. S8). Next, their phosphorylation of GhTIFY3b-mA and GhTIFY3b-mD by GhCPK28 were further examined by in vitro kinase assay. The results showed that GhTIFY3b-mD mimics phosphorylation but can’t be phosphorylated. There is also no phosphorylation in the absence of GhCPK28 in the assay. This means another site was phosphorylated in GhTIFY3b by GhCPK28 (Fig. 6B).

**Figure 6.**
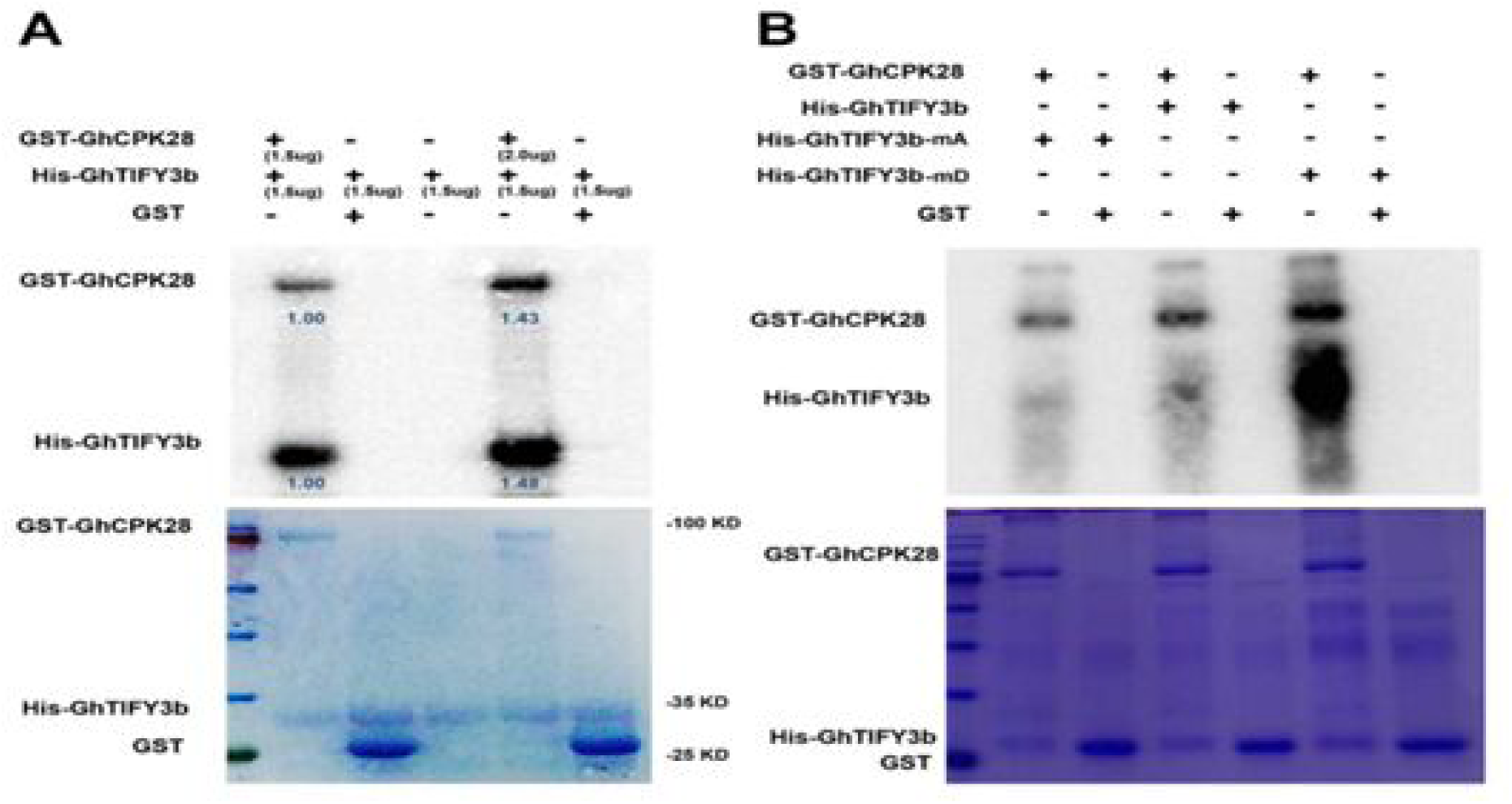
GhCPK28 phosphorylates GhTIFY3b. A, Phosphorylation assays between GhCPK28 and GhTIFY3b in vitro using Phos-tag SDS-PAGE. GST-GhCPK28 was used to phosphorylate purified His-tagged GhTIFY3b protein. A reaction mixture without kinase or without substrate was used as control. The reaction mixture was subjected to SDS-PAGE with Phos-tag, and the phosphorylated protein was immunoblotted with anti-His antibodies. B, In vitro phosphorylation assays of recombinant His-GhTIFY3b, His-GhTIFY3b-mA, or His-GhTIFY3b-mD by GST-GhCPK28 using Phos-tag SDS-PAGE. His-tagged proteins were incubated with immobilized GST or GST-GhCPK28 proteins, and immunoprecipitated portions were detected with anti-His or anti-GST antibodies.

### Overexpression *GhCPK28* decreases the stability of *GhTIFY3b*

To further explore the biological relevance of the phosphorylation of GhTIFY3b by GhCPK28, we used the firefly luciferase (LUC) as a reporter to detect the protein stability of GhTIFY3b. As shown in Figure 7, GhCPK28 was found to have a certain effect on the stability of GhTIFY3b according to the indicated constructs with equal concentrations and volumes were co-infiltrated into *N. benthamiana* leaves by *A. tumefaciens* infiltration. After overexpression of *GhCPK28*, the expression of GhTIFY3b protein decreased and its activity decreased. GhCPK28 does not affect the S-28-mA mutant protein but affects the S-28-mD protein (Fig. 7A). Among 62sk+ GhTIFY3b in the control group, GhTIFY3b-LUC had the highest active enzyme activity, while in 62SK-GhCPK28 + GhTIFY3b in the experimental group, after the addition of protein GhCPK28, the enzyme activity of GhTIFY3b -LUC reporter gene was significantly reduced, indicating that GhCPK28 may degrade the protein of GhTIFY3b-LUC and lead to decreased enzyme activity. There was no change in the enzyme activity of GhTIFY3b-mA after mutation, indicating that GhCPK28 could not affect the enzyme activity of GhTIFY3b-mA-LUC. After mutation at the GhTIFY3b-mD site, the enzyme of GhTIFY3b-mD activity was decreased, indicating that GhCPK28 affected the protein of GhTIFY3b-LUC (Fig. 7B). These data suggested that GhCPK28-mediated phosphorylation reduced the stability of GhTIFY3b.

**Figure 7.**
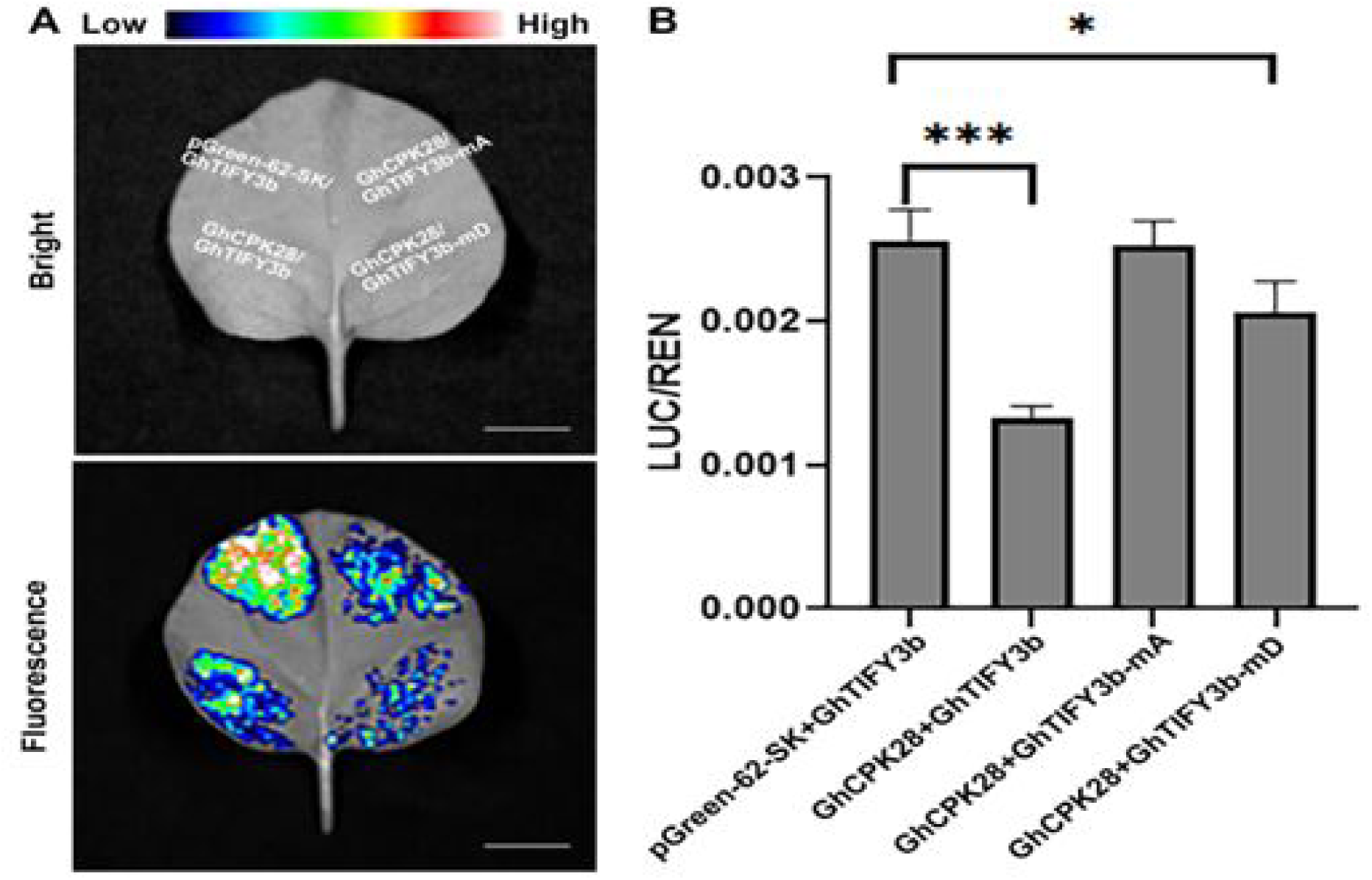
GhCPK28-phosphorylated GhTIFY3b showed decreased protein stability. A, GhTIFY3b, GhTIFY3b-mA, or GhTIFY3b-mD -LUC protein levels with empty vector (EV) or GhCPK28 in *N. benthamiana* leaves. B, The levels of LUC activity of GhTIFY3b, GhTIFY3b-mA, or GhTIFY3b-mD in *N. benthamiana* leaves in the presence of GhCPK28. The LUC/REN activity ratios are means ± sd, n = 3. Bars = 1 cm.

Considering that *GhCPK28* is a negative regulator of Verticillium wilt, and *GhTIFY3b* expression level was up-regulated after *V. dahliae* infection, we also compared the expression of *GhCPK28*-silenced plants before and after *V. dahliae* infection. The expression of GhTIFY3b in *GhCPK28*-silenced plants infected with *V. dahliae* was significantly higher than that in control plants (TRV:00) (Supplemental Fig.S9). These findings indicate that GhCPK28 negatively regulates host defense against *V. dahliae* infection by phosphorylating GhTIFY3b and reduces the stability of GhTIFY3b.

## Discussion

Calcium ions function as a key second messenger in plant immune response (Locato and De Gara, 2018). Calcium-dependent protein kinases (CDPKs or CPKs) are the key proteins of plant signal transduction, transmitting Ca^2+^ messenger through their regulated phosphorylation of various substrates (Liese and Romeis, 2013). Previous study found that CPKs play a pivotal role in plant disease resistance, being either positive or negative regulators (Boudsocq and Sheen, 2013; Romeis and Herde, 2014). In *Arabidopsis*, several CPK proteins are reported to be involved in disease resistance. AtCPK 4/5/6/11 proteins play positive role in disease resistance via phosphorylating the NADPH oxidase RBOHD (Boudsocq et al., 2010; Dubiella et al., 2013; Gao et al., 2013; Kadota et al., 2014). In contrast, AtCPK28 negatively regulates PAMP-triggered early innate immunity by phosphorylating BOTRYTIS-INDUCED KINASE1 (BIK1) (Matschi et al., 2013; Monaghan et al., 2014). In addition, orthologs of *AtCPK28* in rice (*OsCPK4*, *OsCPK18*) and in tobacco (*NaCPK4*, *NaCPK5)* are also shown to be negative regulators of disease resistance (Yang et al., 2012). Here, we identified the cotton *GhCPK28* gene as a negative regulator of resistance to *V. dahliae*. The expression of *GhCPK28* was down-regulated by *V. dahliae* infection, and silencing *GhCPK28* enhanced the resistance of cotton to *V. dahliae*. These data suggest that *CPK28* genes are widely used as negative regulators of immunity across plant species. We also identified a transcription factor *GhTIFY3b* as a positive regulator of resistance to *V. dahliae.* GhTIFY3b is a putative JAZ protein that is involved in JA signaling. Knocking down of GhTIFY3b results in decreased resistance in cotton. This also supports an involvement of JA signaling in *V. dahliae* resistance.

We provided evidence for GhTIFY3b as a putative substrate of GhCPK28. Besides being both involved in disease resistance, these two proteins show direct physical interactions in BIFC and Co-IP assays. In addition, GhCPK28 could directly phosphorylating GhTIFY3b in vitro. These data point GhTIFY3b as a strong candidate substrate of GhCPK28 is involved in JA signaling in disease resistance. Future research should further investigate how phosphorylation affects the stability or activity of GhTIFY3b and test the relevance of this regulation in disease resistance. Knocking down *GhCPK28* increases the induction of the JA related genes *MYC2*, *OPR3*, and *LOX2* by *V. dahliae* infection, suggesting a positive regulation of JA pathway by *GhCPK28*. Interestingly, *GhCPK28* is repressed at 24 hrs after *V. dahliae* infection in cotton, suggesting that *GhCPK28* might be repressed to allow the enhancement of JA signaling and disease resistance. In *Arabidopsis thaliana*, JA signaling plays an important role in resistance to *V. dahliae* (Fradin et al., 2011), and constitutive activation of JA signaling enhanced resistance to *V. dahlia* (Hu et al., 2018). In *Arabidopsis*, the *CPK28* loss-of-function mutant has an elevated expression of JA-responsive genes and increased accumulation of JA and Ile-JA in a developmental stage and tissue specific manner (Matschi et al., 2013). Therefore, the regulation of CPK28 on JA pathway appear to be different in *Arabidopsis* and cotton. The special and temporal dynamics of GhCPK28 could be further studied to determine if they indeed have opposite functions or the apparent difference reflects temporal and special differences.

Although several CPKs are shown to positively or negatively regulate JA synthesis, its underlying mechanism is not clear. Here, we provide evidence that the negative regulation of *GhCPK28* antagonism may be mediated by the JAZ protein *GhTIFY3b*. Previous studies have shown that JAZ protein is associated with disease resistance. For example, *AtJAZ10* is a negative regulator of both JA signaling and the bacterial pathogen *Pseudomonas syringae* symptom development (Demianski et al., 2011). OsJAZ8 is an inhibitor of JA signal via interacting with the JA receptor OsCOI1 and is involved in the resistance to rice bacterial blight (Yamada et al., 2012). *BrTIFY3b* in *Brassica rapa* is induced in expression during *Fusarium oxysporum* infection (Saha et al., 2016). Here we found that knocking down *GhTIFY3b* decreased cotton tolerance to *V. dahliae* infection. GhCPK28 can interact with and phosphorylates GhTIFY3b at Thr-110. These results suggest that GhTIFY3b could be a link between GhCPK28 and JA signaling.

CPK28 proteins regulate multiple pathways which might differ among plant species. Previous work has shown that *AtCPK28* phosphorylates the key immune signaling receptor-like cytoplasmic kinase BOTRYTIS-INDUCED KINASE 1 (BIK1), leading to its degradation by the 26S proteasome and resulting in the attenuation of PTI signaling (Monaghan et al., 2014; Monaghan et al., 2015). In this study, we identified GhTIFY3b as a candidate substrate for *GhCPK28* in regulating disease resistance. Future examination of the role of cotton BIK1 should determine whether or not different substrates and pathways are regulated by CPK28 in *Arabidopsis* and cotton for disease resistance.

In summary, we identified GhCPK28 and GhTIFY3b as regulators of plant immunity. GhCPK28 interacts with and phosphorylates GhTIFY3b, which may influence the JA signaling pathway. Upon infection by *V. dahliae*, *GhCPK28* is repressed, likely contributing to the full induction of JA pathway for disease resistance. These findings not only revealed regulatory components in *V. dahliae* resistance but also offer genetic targets for improving disease resistance in cotton and other crops.

## Materials and methods

### Plant materials and growth conditions

Upland cotton (*Gossypium. hirsutum* L.) TM-1 variety was used in this study. The seeds of TM-1 were sown in pots of soil mixed with vermiculite, and the cotton seedlings were placed in a greenhouse at 28 °C and grew 16-h-light/8-h-dark conditions.

### Pathogen culture and inoculation

The highly aggressive defoliating isolate of *V. dahliae* V991 strain was cultured in Czapek medium (1 g KH_2_PO_4_, 2 g NaNO_3_, 1 g MgSO_4_·7H_2_O, 1 g KCl, 2 mg FeSO_4_·7H_2_O, and 30 g Sucrose L^−1^) at 28 °C. The spores were harvested after 5-d of culture. During infection, sterile distilled water was used to adjust to the spore density to 1 × 10^6^ mL^−1^. The expression of *GhCPK28* was analyzed in 3-week-old cotton seedlings treated with *V. dahliae*, methyl jasmonate, and H_2_O_2_. Cotton seedlings roots were soaked in spore suspension of *V. dahliae* for 30 min and harvested at 0, 0.5, 2, 6, 12, 24, and 48 hours-post-inoculation (hpi) for *V. dahliae* treatment. Cotton seedling roots sprayed with 200 mM methyl jasmonate, 2 mM SA, and 1 mM H_2_O_2_ leaves were collected at 0, 0.5, 1, 2, and 3h, respectively. All samples collected were frozen in liquid nitrogen and preserved at -80 °C for RNA extraction. In parallel, three-week-old seedlings were mock-inoculated with sterile water as control.

### Sequence alignment and phylogenetic analysis

*Arabidopsis thaliana* and upland cotton CPK28 protein sequences were downloaded from https://arabidopsis.org and https://cgp.genomics.org.cn, respectively. ClustalW was used to compare GhCPK28 sequences, and MEGA5 software was used to generate *GhCPK28* phylogenetic tree by the neighbor-joining method (Tamura et al., 2011).

### Total RNA extraction and RT-qPCR

Total RNAs was extracted from 100 mg cotton seedling samples with different treatments by grinding with Plant Total RNA Purification Kit (TIANGEN, China). RNA was then treated with DNase using the DNA-free kit following the manufacturer’s protocol. The cDNA synthesis was performed according to the instructions of the cDNA synthesis SuperMix kit (TransGen biotech, China). RT-qPCR analysis was conducted using SYBR Green Real-time PCR Master mix (Toyobo, Japan) and Real-time PCR detection system (Applied Biosystem Step-One). Cotton *GhUBQ7* acted as internal reference gene. Relative transcript levels using the 2^−ΔΔCT^ or 2^−ΔCT^ methods were calculated (Livak and Schmittgen, 2001). All experiments were performed three times. Primers used for RT-qPCR are provided in Supplemental Table S1.

### Gene cloning, VIGS, and disease assay in cotton

The *A. tumefaciens*-mediated VIGS methods are based on tobacco rattle virus (TRV) vector as previously described (Gao et al., 2011). The gene specific fragments were amplified by PCR using plasmids carrying *GhCPK28* and *GhTIFY3b* cDNA sequences. PCR products were digested with *Kpn* I and *EcoR* I and inserted into the binary *TRV:00* vector, which was digested with same restriction enzymes to form *TRV:GhCPK28* and *TRV:GhTIFY3b* constructs. These plasmids were transformed into *Agrobacterium* strain GV3101. The *Agrobacterium* cultures (OD_600_ = 1) containing two constructs was injected into the fully expanded cotyledon of 1-week-old cotton seedlings with a syringe at a ratio of 1:1 (v/v). At least 20 plants were inoculated with each construct. The *TRV:GhCLA1* construct served as a positive marker for evaluating VIGS efficiency. At 16 d after injection, the expression levels of the target gene were examined, and the successfully silenced plants were subjected to infection by root dipping in the spore suspension of *V. dahliae* V991 for 2 min. The disease index was scored using at least 20 plants per treatment and repeated at least three times. The severity of the disease symptoms on each cotton seedling was scored according to Xu et al.(Xu et al., 2012). The primers for VIGS are listed in Supplemental Table S1.

### Fungal recovery assay and semi-quantification of *V. dahliae* biomass

The same part of the cotton stem segment was harvested 2-week after *V. dahliae* inoculation and sterilized with 10% (v/v) H_2_O_2_. After washing with sterile water for 5 times, the stem sections were placed on potato dextrose agar medium and cultured at 28 °C for 5 days. The number of colonies that the fungus grew on stem segments was considered to be the extent to which the fungus colonizes. The *V. dahliae* biomass was analyzed by semi-quantitative PCR analysis. DNA was extracted from control and *GhCPK28* or *GhTIFY3b*-silenced cotton plants 14 days after *V. dahliae* infection. Fungus-specific forward primer *ITS1-F* and *V. dahliae* specific reverse primer *ST-VE1-R* were applied to amplify the internal transcribed spacer (ITS) region of the ribosomal DNA (Fradin et al., 2011). *GhUBQ7* was used as an internal control. The primers used are listed in Supplemental Table S1.

### Visualization of the levels of H_2_O_2_, and cell death

To visualize the accumulation of H_2_O_2_, leaves of control plants and *GhCPK28* or *GhTIFY3b*-silenced plants were collected 24 hours after *V. dahliae* inoculated. The leaves were incubated in DAB solution (1 mg/mL, pH 3.8) for 8 hours, decolorized with 95% ethanol for 2min, and then decolorized with absolute ethanol until the green color of the leaves was completely removed. The fading process was further faded in a boiling water bath. The accumulation of H_2_O_2_ in 70% glycerol was observed under microscope.

The plant cell death was observed by Trypan blue staining. The leaves were soaked in Trypan blue dye (1 mg trypan blue, 1 g phenol, 1 ml lactic acid, and 1 ml glycerol dissolved in 1 ml sterile distilled water), then boiled and stained. After cooling to room temperature, the sample was decolorized with chloral hydrate solution (2.5 g/ml). Each experiment was repeated three times.

### Measurement of H_2_O_2_ and NO content

Three plants leave of *TRV:00*, *TRV:GhCPK28* and *TRV:GhTIFY3b* were randomly collected every 10 min interval after *V. dahliae* inoculation 0 to 60 min. The contents of hydrogen peroxide (H_2_O_2_) and NO were determined with a Quantitative Assay Kit (Jiancheng, China). Three biological experiments which consisted of 3 plants per experiment were measured.

### Y2H assay

The yeast (*Saccharomyces cerevisiae*) two-hybrid assay was conducted according to the manufacturer’s instructions for the Match-maker Gold Yeast Two-Hybrid System (Clontech, USA). To generate the bait construct BD-GhCPK28, the coding sequence (CDS) of *GhCPK28* was amplified and cloned into the pGBKT7 vector. As to generate the AD-GhTIFY3b, the coding sequence (CDS) of *GhCPK28* was amplified and cloned into the pGADT7 vector. BD-*GhCPK28* and AD-GhTIFY3b were co-transferred into the yeast strain AH109 for screening a library prepared from cotton roots under *V. dahliae* treatment conditions. Yeast strain AH109 cells co-transformed with these constructs were inoculated on SD-Leu-Trp, SD-Leu-Trp-His (with X-α-Gal), and SD-Leu-Trp-His-Ade (with X-α-Gal) media. The primers for Y2H assay are shown in Supplemental Table S1.

### Recombinant protein expression and pull-down assays

The full length of *GhCPK28* was amplified and cloned into the pGEX-4T-1 (Pharmacia) to generate the GST-GhCPK28 protein and the full length of *GhTIFY3b* was amplified and cloned into the pCzn1vector (Zoonbio) to generate the His-GhTIFY3b proteins. The production of GST and His fusion recombinant protein from *Escherichia coli* strain BL21 was induced by isopropyl β-D-1-thiogalactoside (IPTG). After adding IPTG, the protein was induced overnight at 15 °C. The soluble GST and His fusion proteins were purified using the MagneGST Protein Purification System (Promega, USA) or the MagneHis Protein Purification System (Promega) according to the manufacturer’s instructions. Purified mixtures of pCzn1 and pGEX-4T-1 without inserted transgenic were used as negative controls for the His-tagged fusion protein and GST-tagged fusion protein, respectively. The pull-down experiment between GST-GhCPK28 and His-GhTIFY3b was performed as previously reported (Deng et al., 2017). The primers used are listed in Supplemental Table S1.

### BiFC assays and subcellular localization

To generate the BiFC constructs, the full lengths of *GhCPK28* and *GhTIFY3b* were amplified with primer pairs of *GhCPK28* and *GhTIFY3b* and inserted into linearized pCAMBIA1300-35S-NY173 or pCAMBIA1300-35S-YC155 vectors to obtain GhTIFY3b-nYFP and GhCPK28-cYFP. To determine the subcellular localization of *GhCPK28* and *GhTIFY3b*, the full lengths of these genes were amplified with corresponding primer pairs and cloned into vector pCAMBIAl302 to generate the C-terminally fused YFP constructs. Proteins fused with cYFP and nYFP were co-expressed via Agrobacterium strain GV3101 infiltration in *N. benthamiana* (Chen et al., 2008). The fluorescence signals of leaf epidermal cells were observed by confocal microscope (Nikon C2-ER). The primers used are listed in Supplemental Table S1.

### In vitro kinase assay

One μg of purified recombinant GST-GhCPK28 fusion protein (expressed from *E. coli*) was incubated with 5-10 μg purified recombinant His-tagged substrate in a kinase reaction buffer (20 mM Tris-HCl pH 8.0, 20 mM MgCl_2_, 1 mM DTT, 50 mM ATP) with or without 1 μCi [γ-^32^P] ATP at 30 °C for 30 min. Proteins from the reaction were separated by SDS-PAGE. Coomassie blue was used to stain gels of proteins (the reaction without [γ-^32^P]). Autoradiography was used to detect phosphorylation signal from reactions with [γ-^32^P]. GhTIFY3b-mD simulating the continuous phosphorylated state and GhTIFY3b-mA simulating the non-phosphorylated state.

### Co-IP Assay

The full lengths of GhCPK28 and GhTIFY3b were amplified by primer pairs and inserted into pBWA(V)HS-3xflag-2 and pBWA(V)Hs-TMV-MYC vectors. GhCPK28-flag and GhTIFY3b-MYC constructs containing C-terminally tagged flag protein and MYC protein were generated, respectively. All plasmids were transformed into *A. tumefaciens* strain GV3101. The clones carrying GhCPK28-flag and GhTIFY3b-MYC, as well as the clones carrying the control vector, were coinfiltrated into the leaves of *N. benthamiana* through *A. tumefaciens* infiltration (Chen et al., 2008). The infiltrated plants were grown at 25°C for 48 to 72 h, and the tissue of infiltrated leaves was collected and frozen with liquid nitrogen. About 1 g of infiltrated leaf tissue was ground into a powder in liquid nitrogen and homogenized in 600µl of IP Lysis buffer containing a protease inhibitor cocktail (1x). Follow the instructions for protein A/G beads and western blot tests to perform immunoblotting and Co-IP experiments to visualize the results. The primers used in this study are listed in Supplemental Table S1.

### Luciferase Reporter Assays in *N. benthamiana*

The full lengths of native GhTIFY3b, phosphodeficient GhTIFY3b-mA, and phosphomimic GhTIFY3b-mD were amplified with primer pair GhTIFY3b-pGreenII-0800-F/R using the GhTIFY3b-pET-28a, GhTIFY3b-mA-pET-28a, and GhTIFY3b-mD-pET-28a plasmids as templates, respectively. The PCR-amplified fragment was inserted into a linearized pGreenII-0800 vector digested with *EcoR*I and *Nco*I. GhTIFY3b-pGreenII-0800, GhTIFY3b-mA-pGreenII-0800, and GhTIFY3b-mD-pGreenII-0800 vectors were generated by in-fusion enzyme to obtain dual-luciferase reporter system. The full length of GhCPK28 was amplified by primer pairs of GhCPK28-pGreenII-62-SK-F/R and inserted into the linearized pGreenII-62-SK vector digested by *Bsa*I and *EcoR*I. pGreenII-62-SK vector without DNA insertion served as empty vector control. All the plasmids were transferred into *A. tumefaciens* strain GV3101, an transformed into *N. benthamiana* for transient expression. The *A. tumefaciens* containing GhTIFY3b-pGreenII-0800, GhTIFY3b-mA-pGreenII-0800 and GhTIFY3b-mD-pGreenII-0800 vectors were adjusted to the same concentration of OD600 = 0.2 and the empty vector and GhCPK28-pGreenII-62-SK were adjusted to OD600 = 0.6. These corresponding coinfiltrated constructs were mixed in equal volumes and coinfiltrated into *N. benthamiana* leaves to ensure comparability. The infiltrated plants grew at 25°C for 2 days. The back of infiltrated leaves was sprayed with the solution of luciferin potassium salt (0.2mg/ml) and then leave it in the dark for 5-10min to take photos. The LUC luminescence was observed with a cryogenically cooled CCD camera (Lumazome PyLoN 2048B) and the exposure time was set to 15 min. The primers used in this study are listed in Supplemental Table S1.

## Accession numbers

The sequence data for this article can be found in the CottonFGD database (https://cottonfgd.org) or GenBank databases with the following accession numbers: GhCPK28 (six copies in the whole genome), Gh_A02G1635, Gh_D03G0087, Gh_A10G0886, Gh_D10G0863, Gh_D11G1774 and Gh_A11G161; GhTIFY3b, LOC107907061 or Gh_D05G039400.1; and GhUB7, DQ116441.

## Author contributions

S.G., W.S., J.W., Z.Z., W.H. and S.P. performed the experiments. B.Z., J.H. and Q.H. conceived and designed the research. S.G., W.S., W.J., Z.Z., W.H. and S.P. analyzed the data. Q.H. and J.H. supervised the work. B.Z. and Q.H. wrote the article and J.H. contributed and supervised the final article version. B.Z. and Q.H. agree to serve as the author responsible for contact and ensure communication. All authors read and approved the manuscript.

## Supplemental data

The following materials are available in the online version of this article.

---

**Supplemental Figure S1.** Characterization of *GhCPK28*. A, Diagram of the *GhCPK28* gene structure. The green box indicates the EF-hand domains, and the blue frame indicates conserved amino acids. B, Phylogenetic analysis of *GhCPK28* homologues from *Oryza sativa* (Os), *Zea mays* (Zm), *Gossypium raimondii* (Gr), *Solanum lycopersicum* (Sl), *Arabidopsis thaliana* (At) and *Gossypium hirsutum* (Gh). The neighbor-joining tree was built using the MEGA5 program.

**Supplemental Figure S2.** Amino acid sequence alignment of GhTIFY3b and AtTIFY3b.

**Supplemental Figure S3.** Subcellular localization of GhCPK28 and GhTIFY3b. Localization of transient expression of GhCPK28-GFP and GhTIFY3b-GFP in *N. benthamiana* leaves. Co-localization of GhCPK28-GFP in the cell membrane and GhTIFY3b-mCherry in the nucleus. Scale bar = 50 μm.

**Supplemental Figure S4.** Expression profile of *GhTIFY3b*. A, RT-qPCR analysis of *GhTIFY3b* expression at different time points after inoculation with *V. dahliae*. Total RNA was extracted from roots of control and *V. dahliae*-inoculated plants. Error bars represent ± SD (n=3). The standard deviations were calculated from the results of three independent experiments (** P< 0.01, *** P< 0.001, t-test). B, RT-qPCR analysis of *GhTIFY3b* expression in JA-, SA- and H_2_O_2_-treated cotton. Total RNA was extracted from roots of control and JA-, SA- or H_2_O_2_-treated plants. The values are means ± sd, n = 3, and normalized to those of *GhUBQ7*. Different letters indicate significant differences at P< 0.05 (ANOVA’s multiple comparison test).

**Supplemental Figure S5.** Amino acid alignment of six homologous *TIFY3b* genes from upland cotton.

**Supplemental Figure S6.** Silencing of *GhTIFY3b* results in ROS elimination and cell death. A, ROS and cell death in of *TRV:00* and *TRV:GhTIFY3b* plant leaves were observed by DAB-peroxidase staining and cell death after 24 h inoculation with *V. dahlia*. B, NO content in roots of *TRV:00* and *TRV:GhTIFY3b* plants 1 h after inoculation with *V. dahliae.* Error bars represent SD of three technical replicates of one biological experiment. The standard deviations were calculated from the results of three independent experiments (* P< 0.05, ** P< 0.01, t-test).

**Supplemental Figure S7.** Predicted GhTIFY3b phosphorylation sites. GhTIFY3b T110 is phosphorylated by GhCPK28 with MS analysis. The confident b2 and b3 ions provide strong evidence for phosphorylation of the third Thr residue.

**Supplemental Figure S8.** GST pull-down assay showed that GST-GhCPK28 had direct interaction with His-GhTIFY3b-mA or His-GhTIFY3b-mD fusion protein. His-tagged proteins were incubated with immobilized GST or GST-GhCPK28 proteins, and immunoprecipitated fractions were detected with anti-His or anti-GST antibodies.

**Supplemental Figure S9.** Silencing *GhCPK28* leads to increase expression of *GhTIFY3b*. qRT-PCR analysis of *GhTIFY3b* expression level in *TRV:00* and *TRV:GhCPK28* plants inoculated with *V. dahliae*. Error bars represent SD of three technical replicates of one biological experiment. The standard deviations were calculated from the results of three independent experiments (* P< 0.05, ** P< 0.01, t-test)

**Supplemental Table S1.** Primers used for this study.

**Supplemental Table 2.** The potential GhCPK28 interaction proteins in the Y2H screening.

